# Electrodiffusion model of synaptic potentials in dendritic spines

**DOI:** 10.1101/274373

**Authors:** Thibault Lagache, Krishna Jayant, Rafael Yuste

## Abstract

When modeling electric current flow in neurons and excitable cells, traditional cable theory ignores electrodiffusion (i.e. the interaction between electric fields and ionic diffusion) as it assumes that concentration changes associated with ionic currents are negligible. This assumption, while true for large neuronal compartments, fails when applied to femto-liter size compartments such as dendritic spines - small protrusions that form the main site of synaptic inputs in the brain. Here, we use the Poisson (P) and Nernst-Planck (NP) equations, which relate electric field to charge and couple Fick’s law of diffusion to the electric field, to model ion concentration dynamics in dendritic spines. We use experimentally measured voltage transients from spines with nanoelectrodes to explore these dynamics with realistic parameters. We find that (i) passive diffusion and electrodiffusion jointly affect the kinetics of spine excitatory post-synaptic potentials (EPSPs); (ii) spine geometry plays a key role in shaping EPSPs; and, (iii) the spine-neck resistance dynamically decreases during EPSPs, leading to short-term synaptic facilitation. Our formulation can be easily adopted to model ionic biophysics in a variety of nanoscale bio-compartments.

## INTRODUCTION

Dendritic spines, nanoscale bulbous protrusions located along dendrites of principal neurons, form the primary site of excitatory synaptic input in the mammalian brain [1–3]. Their function and plasticity are essential for neuronal function, memory formation and brain development [4, 5]. Excitatory neurotransmitters released during synaptic transmission activate receptors on the spine head, which causes ion channels to open and current to flow into the spine. This current charges the spine causing an excitatory post-synaptic potential (EPSP), which subsequently integrates onto the dendrite and summates with other synaptic inputs on its way to the soma and axon initial segment [6]. When this summation crosses a threshold, the neuron fires an action potential (AP). Spines thus present the first node in the path of neuronal integration, and voltage dynamics (i.e. rise time, fall time and amplitude) within a spine during synaptic input determine the characteristics of the downstream signal. Voltage recordings from this nanoscale neuronal subdomain have been challenging, and measurements have traditionally been at odds with each other, specifically with regard to the spine head EPSP magnitude [7–12] and neck resistance values [7, 8, 12–16]. Recently however, optical and electrical recordings have reported fast and large EPSP values in the spine with high neck resistances [8, 10, 15, 17]. But, given that spines exhibit a stereotypical, but highly variable geometry [18] comprising a bulbous head (volume ranges ~ 0.01-0.3 μm^3^) connected to the parent dendrite across a narrow neck (length ranges ~ 0.1-5 μm; diameter ranges ~ 5-200 nm), there is a need for models to explore the relationship between geometry and voltage dynamics during synaptic inputs. Such models may help reconcile the different values that have been experimentally procured and possibly help shed intuition on the reasons for the discrepancies in literature.

Traditionally, spine biophysics and the EPSP map across a neuron are modeled using cable theory (CT) [19–23]. The cable equations treat the membrane as an RC circuit and compute the time evolution of the signal usually in the form of partial differential equations. This formulation neglects the effect of concentration gradients, which is a reasonable hypothesis for large neuronal compartments (such as the original squid giant axon) but may fail to accurately describe electrostatics in nanoscale structures such as dendritic spines. In addition, neck resistance has been indirectly estimated with fluorescence recovery after photo-bleaching (FRAP) time constants [13, 14, 24, 25] (i.e. charged or uncharged dye molecules diffusing back into the spine head upon a spine head photo-bleaching). Here, the mean escape time τe of a single diffusive molecule (with diffusion coefficient D) from the spine head into the dendrite can be described by the relation[26] 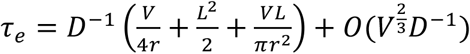, where V is the volume of the spine head, r the radius of the cylindrical neck and L its length. For a typical spine with head volume V = 0.1 μm^3^, neck length L = 1 μm, neck radius r = 50 nm and a diffusion constant D = 0.5 × 10^3^ μm^2^ s^-1^ (corresponding to the measured coefficient of sodium ions in cytoplasm) [27, 28], we obtain that 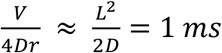 and 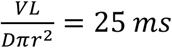. Thus, in most spine geometries we can approximate 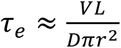, which is then used to fit the exponential decay rate of the FRAP transient. Neck resistance is finally estimated with 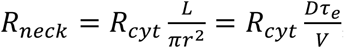, where *R_cyt_*, the longitudinal cytoplasmic resistivity, is assumed to be constant ≈ 1 – 1.5 Ω. m [11, 22]

While both cable theory and FRAP measurements have largely been used to model and extract spine biophysical parameters [7, 14, 21, 25], they do have limitations and many of the values are at odds with many experiments [9, 11, 24]. As previously mentioned, one critical aspect that is ignored in cable theory is electrodiffusion [29–32] (i.e. the effect of the electric field on the concentration gradient), which is important when large longitudinal voltage gradients and concentration changes occur. This aspect becomes more critical in the small dendritic spines where sizeable voltage swings, large concentration changes on the order ~mM, and appreciable electric field gradients across a nanoscale neck can occur within a few milliseconds. Such effects could modulate both voltage and current transmission at a fundamental level.

In order to accurately describe electrolyte dynamics in biological micro-domains and explicitly account for the effect of geometry and electrodiffusion, the Poisson-Nernst-Planck (PNP) formalism is ideal [33]. However, the PNP equations cannot be solved analytically in complex three-dimensional structures, such as dendritic spines. Previous work modelling electrodiffusion effect in spines [29, 34] have either captured the effects of the PNP purely through numerical simulations [12], examined dynamics under non-electroneutral conditions [12], or solved PNP equations only in cylindrical geometries [29]. Here, we use singular perturbation theory and derive coarse-grained PNP equations to describe voltage dynamics in the spine head during synaptic input, explicitly accounting for the effect of geometry and electrodiffusion. We find that electrodiffusion can play a significant role in determining the overall EPSP magnitude and time scales. Moreover, and in contrast to cable theory analysis, we show that electrodiffusion predicts current modulation across the neck on a millisecond time scale, which suggests that the neck resistance can dynamically vary as a function of synaptic current. We use recently published measurements of spontaneous EPSP transients from spines to highlight these effects under different geometries.

## RESULTS

### PNP formalism

We used singular perturbation theory and analyzed the dynamics of both positive and negative charges inside the spine head and neck (Fig. 1a), and derived a novel coarse-grained system of equations that fully captures the coupled dynamics of ions and voltage with the PNP formalism. It uses a coupled system of two differential equations to describe the interacting ion and electrical potential dynamics: (1) the Poisson (P) equation (Eq. 1) that computes the electrical potential rising from the local differences between the concentrations of negative *c^-^(x, t)* and positive *c^+^(x; t)* charges,

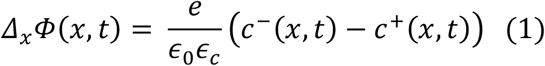

**Figure 1:**
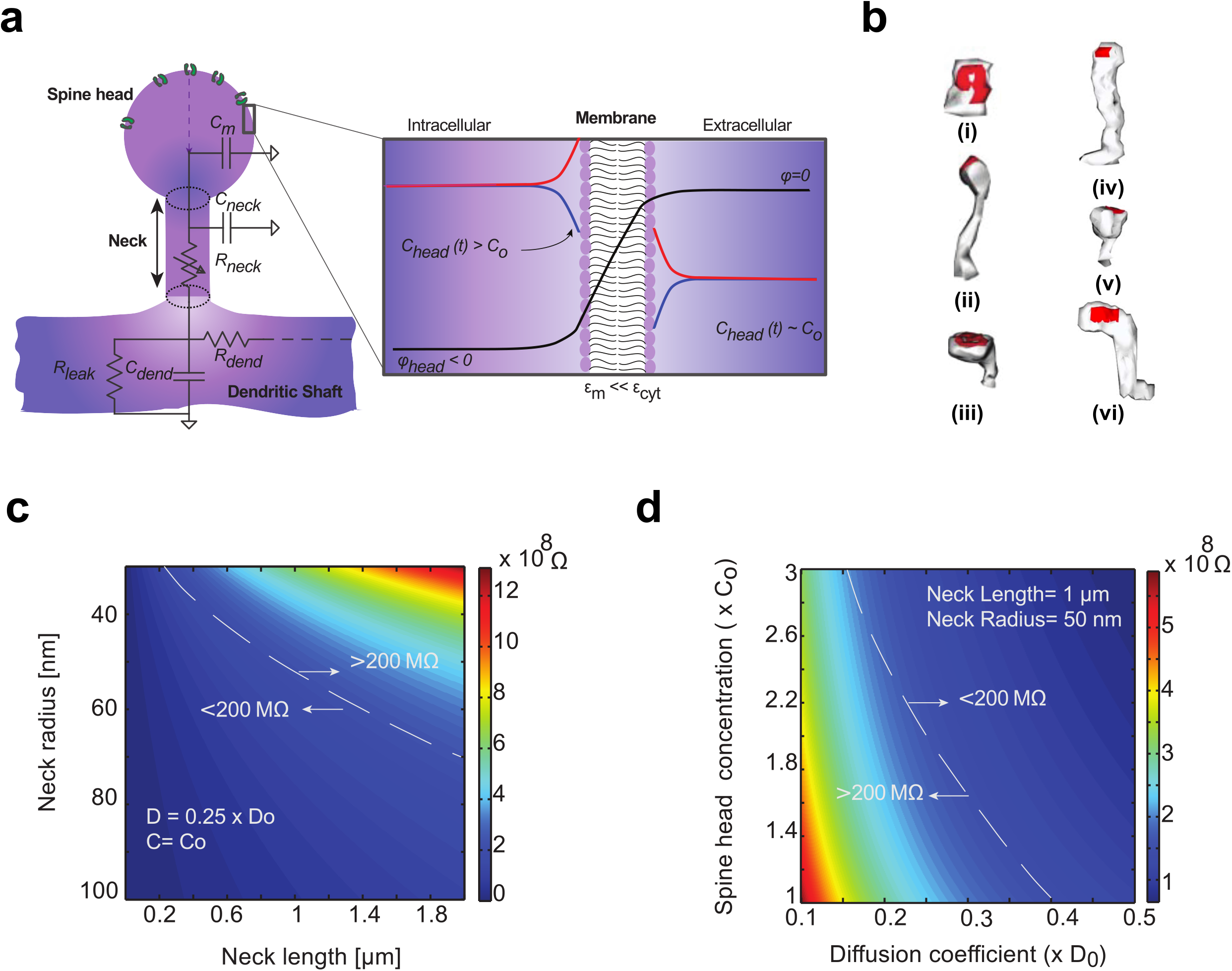
Electro-diffusion model of dendritic spine. **a**) Spine electrical geometry. Spine head contains synaptic post-synaptic density and ion channels. Membrane is modeled as an impermeable dielectric with small electrical permittivity compared to cytoplasm (ϵ_m_ ≪ ϵ_c_). At resting state, the spine head is depolarized with negative electrical potential Φ_0_ ≈ –60 mV. Ion concentration c_head_(t) varies compared to bulk concentration c_Q_. Thin cylindrical neck links the spine head to the parent dendrite. Equivalent circuit describing the dendritic spine electrostatics is represented. Here Cm denotes the capacitance of the head membrane, Cneck and Rneck denote the membrane capacitance and the longitudinal cytoplasmic resistance of the spine neck. Cdend and Rdend denote the membrane and resistance of the dendrite. An additional membrane resistance Rleak models the ion leaks through the dendrite membrane. All of which combine to determine the voltage kinetics inside the spine head. **b**) Ultra-structural images of dendritic spine morphology (data reproducedfrom [12]). Red represents postsynaptic densities **c**) Geometrical determinants of passive spine neck resistance. **d**) Physiological determinants of spine neck resistance.

Where, 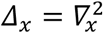 is the Laplacian operator, *e* the elementary electrical charge, *ε*_0_ the vacuum permittivity and *ε_c_* the relative permittivity of the cytoplasm, and (2), the Nernst-Planck (NP) equation (Eq. 2) which captures the contribution of the electrical field on the concentration gradient;

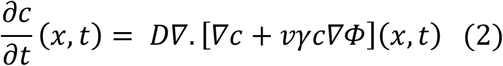

Where, *D* is the diffusion constant (that we assumed to be the same for both positive and negative charges), 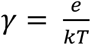, and *v* = ±1, is the ionic valence. An analytical solution to the full system of PNP equations is not possible and hence must be either numerically computed or asymptotically estimated.

### Modeling electrostatics in the spine head and neck

We approximated the geometry of the dendritic spine with a ball (spine head, radius R) connected to the parent dendrite across a cylindrical thin neck (length L, cross-section *S* = *πa*^2^, with *a* the neck radius that we assumed to be constant (Fig. 1a). We also assumed that the neck radius is much smaller than the head radius *a* ≪ *R*, as corroborated by ultra-structural reconstructions [18] (Fig. 1b) and super-resolution microscopy of living spines[14]. We modeled the spine head membrane as an impermeable membrane (no ion leak), with thickness d and small electrical permittivity *ϵ_m_* ≪ *ϵ_c_* (Fig. 1a, inset) (Table 1). We also assumed the continuity of the electrical field (derivative of the electrical potential) at the membrane boundary, a boundary layer condition employed to solve electrostatic equations across dielectric layers [35, 36]. Finally, boundary conditions for the potential and ion concentrations at the neck entrance were matched to physiological solutions inside the spine neck.

**Table 1.** Parameters of the electrodiffusion model.

| Variable | Name | Value | Reference |
| --- | --- | --- | --- |
| $e$ | Elementary charge | $\approx 1.6 \cdot 10^{-19} \text{C}$ | |
| $\epsilon_0$ | Vacuum permittivity | $\approx 8.85 \cdot 10^{-12} \text{Fm}^{-1}$ | |
| $\epsilon_c$ | Cytoplasmic permittivity | 80 (= water) | |
| $\epsilon_m$ | Membrane permittivity | 2 (= phospholipid bilayer) | |
| $\gamma = \frac{e}{k_B T}$ | NP constant | $\approx 37 \text{V}^{-1}$ | |
| $R$ | Radius of the spine head | 50 – 500 nm | [12] |
| $L$ | Length of the spine neck | 0.1 – 2 $\mu\text{m}$ | [12] |
| $a$ | Neck radius | 30 – 100 nm | [12] |
| $S = \pi a^2$ | Cross section of the spine neck | $0.28 - 3.1 \cdot 10^{-2} \mu\text{m}^2$ | |
| $D$ | Diffusion constant of ions | $0.5 \cdot 10^{-9} \text{m}^2 \text{s}^{-1}$ | [19-20] |
| $g_0$ | Maximum synaptic conductance | 1.0 – 8.7 nS | Fitted to data |
| $c_m$ | Membrane capacitance per unit of surface | $0.01 \text{Fm}^{-2}$ | [31] |
| $c_0$ | Bulk ion concentration | 150 mM | [31] |
| $\Phi_0$ | Resting membrane potential | –60 mV | Fitted to data |
| $\mu$ | Kinetics parameter (1) of ion channel opening (sigmoidal function) | 0.27 – 0.71 ms | Fitted to data |
| $\tau_1$ | Kinetics parameter (2) of ion channel opening (sigmoidal function) | 0.075 – 0.2018 ms | Fitted to data |
| $\tau_2$ | Kinetics parameter of ion channel closure (mono-exponential) | 3.756 – 45.54 ms | Fitted to data |
| $\lambda_D = \sqrt{\frac{\epsilon_0 \epsilon_c}{2e\gamma c_0}}$ | Debye length | $\approx 1 \text{nm}$ | |
| $\delta_1 = \left(\frac{\lambda_D}{R}\right)^2$ | Singular perturbation parameter in the spine head | $= 0.02 - 0.002$ | |
| $\delta_2 = \left(\frac{\lambda_D}{L}\right)^2$ | Singular perturbation parameter in the spine neck | $= 0.01 - 0.001$ | |

To mathematically analyze the PNP equations within the spine head, we applied the singular perturbation method using dimensionless variables, and approximate solutions (i.e. asymptotic expansions). Specifically, the dimensionless system of variables is given by 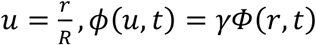, 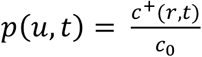 and 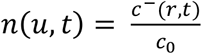, with 0 ≤ *r* ≤ *R* the radial coordinate within the spine head and *c_0_* the total concentration of positive (negative) ions inside the spine cytoplasm at resting state (Table 1). The reduced Poisson equation (Eq. 3) then read (see SI-II-A for details),

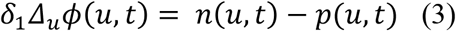

where Δ_*u*_ is the Laplace operator in dimensionless variable u. Here 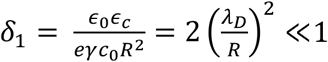, with *λ*_*D*_ the Debye length (*λ*_*D*_ ≈ 1 nm (Table 1)). As *δ_1_* tend to *0*, we obtain electroneutrality in the spine with constant concentration *c*^+^(*r,t*) ≈ *c*^−^(*r,t*) ≈ *c_head_*(*t*), and constant potential *Φ*(*r, t*) ≈ *Φ_head_*(*t*) (Fig. 1a).

This constant potential approximation in the bulk was also confirmed in recent numerical simulations using finite element, steady-state simulations [12], and is true for the entire spine microdomain, barring a thin boundary layer near the membrane- i.e. the Debye layer (Fig. 1a, inset). To obtain the full solution for the potential and the ion concentrations inside the entire spine head domain (i.e. bulk and boundary layer), we compute the inner solutions near the membrane that both match the boundary conditions and the asymptotic solutions in the bulk. We find that, due to the small electrical permittivity of the membrane, the maximal potential drop occurs across the membrane bilayer. The computed bulk potential is then given by (see SI-II-A for details),

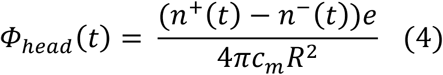

Here *n*^+^(*t*) and *n*^−^(*t*) are the total number of positive and negative ions inside the spine head at time *t*, and ***c_m_*** is the membrane capacitance per unit of surface. Typically, *c_m_* ≈ 0.01 Fm^−2^ (Table 1), and therefore small differences between the total number of positive and negative charges inside the head lead to significant changes of the spine head potential. For example, inside a spherical spine head with radius R = 500 nm, the resting potential *Φ*_0_ ≈ –60 mV corresponds to a net excess of ≈ 12,000 negative charges, while the total number of ions is equal to 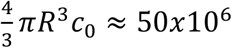 ions. To put this in context, a typical ion channel can flux several thousand ions per millisecond[37]. Thus, due to the small capacitance of the spine head, the entry of relatively few positive charges during synaptic input will result in a rapid depolarization.

The kinetics of ion concentration and potential inside the spine head critically depends on the ionic fluxes with the parent dendrite across the neck. To analyze these fluxes, we followed the methodology developed for modeling ion channels [38, 39] and reduced PNP equations to one-dimensional equations along the neck’s principal axis from the spine head (l = 0) to the dendritic shaft (l = L) (Fig. 1a, see SI-II-B-1 for details). For positive and negative ions, the total flux *J_total_*(*t, v*), with *v* = ±1 (the valency) comprises of a diffusion *J_neck_*(*t*) and current *I_neck_*(*t*) (the drift component)

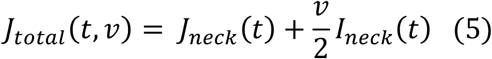

Under electro-neutral conditions, the diffusion flux drives an equal amount of positive and negative charges in the same direction, and thus results in no net electrical current. Mathematical analysis of PNP equations inside the neck leads to (see SI-II-B-2 for details)

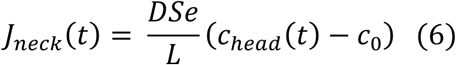

and,

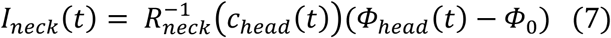

with neck resistance

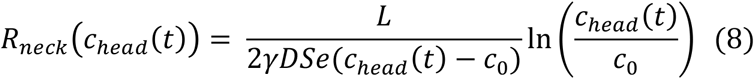

Here, we highlight the following findings: First, the asymptotic diffusion flux (Eq. 6) is the expression normally used to interpret FRAP experiments. This formulation captures only diffusion and neglects the contribution of the electric field on ion dynamics. Second, the neck resistance is not solely dependent on geometry as assumed in cable theory analysis (Fig. 1c) and described in Eq. 9, which depicts the neck resistance assuming constant ion concentration,

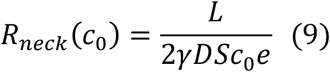

but also critically depends on the ion concentration inside the spine head (Fig. 1d and Eq. 8).

### Fast electrical and slow diffusion dynamics inside the dendritic spine

During synaptic input, both AMPA and NMDA receptors are activated. For the sake of computational simplicity, we chose to neglect NMDA receptors given that AMPA receptors are known to be the major source of Na^+^ current. Moreover, by considering the relative permeability of AMPA receptors to main ions and Goldman-Hodgkin-Katz equation, we computed that the reversal potential of AMPA receptors is 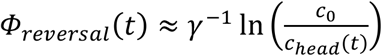 (see SI-II-C-1 for details), and that the synaptic current is governed by the Nernst–Planck equation of transport:

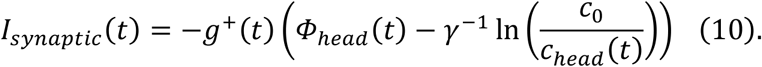

Here *g*^+^(*t*) is the time-dependent conductance of AMPA channels. The ionic influx is thus maximum at resting potential *Φ*_0_ ≈ −60 mV and concentration *c*_0_ and collapsed when the spine head potential and concentration increase.

Using Eq. 6–8 together with the synaptic current (Eq. 10), we obtained the coarsegrained system of differential equations for (see SI-II-C-1 for details)

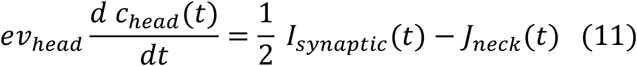

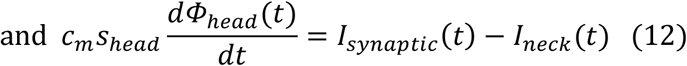

Which we solved self-consistently. Here 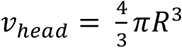 and *s_head_* = 4*πR*^2^ are the volume and the surface area of the spine head respectively. Equation (11) describes the gradual increase of ion concentration inside the spine head during channel opening, while equation (12) captures the potential dynamics. Equations (11) and (12) are non-trivially coupled because synaptic and neck current depends on spine head potential and concentration.

The coarse-grained system of equations (12–13) is a slow-fast dynamical system. The time constant of concentration changes 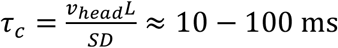 is much longer than the time scale of voltage transients, which is due to the charging/discharging of the spine head (capacitor) through the neck resistance (time constant 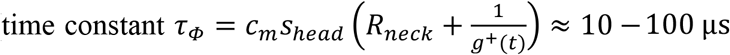) (Supplementary Fig. 1). Thus, for a Heaviside step input (constant channel conductance, *g*^+^(*t*) = *g*^+^), voltage increase is rapid at the onset of ion-channel opening, with significant depolarization values and reaches a plateau given by 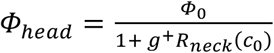 in less than 100 μs (Fig. 2a). We termed the first few hundred microseconds of the transient as the *electrostatic* phase and the millisecond time scale the *diffusion* phase. Concentration changes during the electrostatic phase are negligible, but become significant during the diffusion phase (Fig. 2b). Increased ion concentration has two main effects: the decrease of effective neck resistance (Eq. 8 and Fig. 2c) and reversal potential (Fig. 2d). The rapid voltage depolarization during the electrostatic phase causes a decrease in synaptic current (Eq. 10 and Fig. 2e), mirrored by an increase in neck current (Fig. 2f) which plateaus as voltage reaches a steady state. Synaptic and neck currents are then equal, and subsequently decreases with concentration, as reversal potential decreases and reduces the synaptic electromotive force. We stress that steady state voltage and net ion concentration depend solely on spine neck geometry, whereas their kinetics is controlled by the size of the spine head, volume and membrane capacitance.

**Figure 2:**
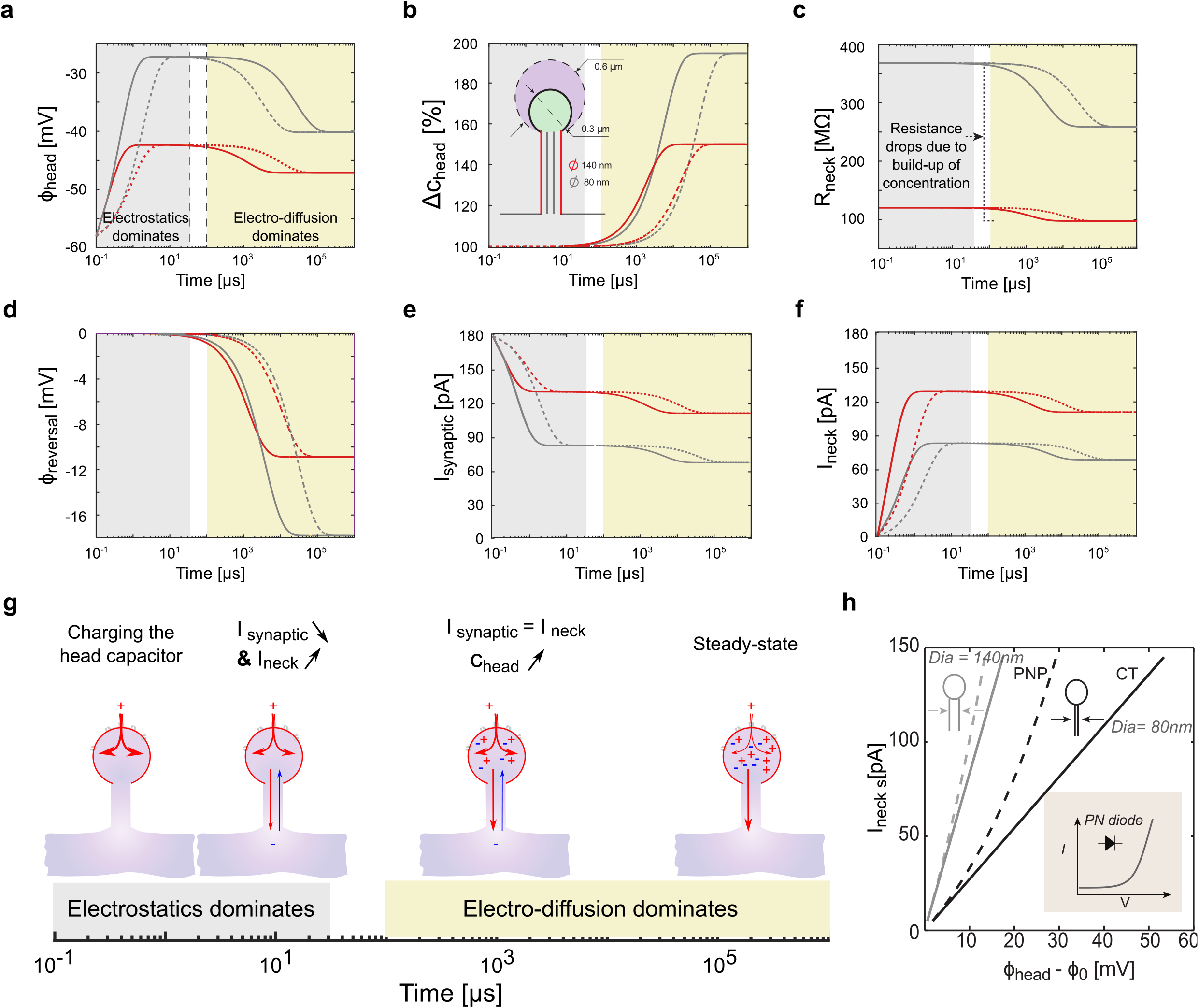
Ionic and voltage dynamics in model. Slow-fast kinetics of ion concentration and electrical potential within the spine head during a step entry of positive ions (constant synaptic conductance g^+^(t) = Constant = 3 nS). **a**) Log plot of the voltage kinetics within different spine geometries, with large (R = 600 nm, dashed line) and small (R= 300 nm, solid lines) head, and different neck diameters (140 nm (red) and 80 nm (grey)). The electrostatics forces dominate at small time scales (< 100 μs) and diffusion at larger time scales (> 1 ms). Note the deflection of the electrical potential due to concentration changes and diffusion. **b**) Log plot of the concentration kinetics. **c**) Log plot of the neck resistance kinetics. Notice the drop in neck resistance as ion concentration increases. **d**) Log plot of the synaptic reversal potential kinetics. **e**) Log plot of the synaptic current kinetics. Note the deflection due to concentration builds-up and decrease of the reversal potential. **f**) Log plot of the neck current kinetics. **g**) Biophysical model: In first microseconds, positive ions flow through AMPA receptors and charge the spine head capacitor, rising head potential. This drives an electrical current though the spine neck (exchange of positive ions with incoming negative ions from dendrite reservoir), and also decreases the synaptic electromotive force and current. A part of positive ions that enter through AMPA receptors accumulate inside the spine head, the global electro-neutrality being conserved thanks to incoming negative ions. Slow increase of ion concentration inside the spine head leads to neck resistance and reversal potential decrease. Concentration reaches a plateau after tens of milliseconds, when concentration gradient of negative charges equilibrates neck current. **h**) I-V curve of dendritic spine for two different neck diameter (140 nm (grey) and 80 nm (black)). The discrepancy between the spine I-V curve (dashed line) and Ohm’s law with constant resistance (solid line) is highlighted. For thin and highly resistive neck, the spine behaves like a diode (inset scheme).

At steady-state, the amplitude of current through the neck *I_neck_*(∞) is equal to the amplitude of the synaptic current *I_synaptic_*(∞) and also to the diffusive outflux *I_neck_*(^∞^), where ∞ denotes the steady state value for *t* ≫ *τ_c_*. These relations lead to an implicit equation for the steady state current that is solved numerically (see SI-II-C-2 for details), and I-V relationship across the spine neck (Fig. 2h and Supplementary Fig. 2)

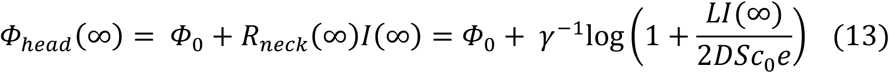

Eq. 13 shows an highly non-linear relation, similar to current rectification observed in nanofluidic diodes[40] (Fig. 2h, inset). Such a non-linear relationship - a consequence of the PNP, was also observed with finite element simulations [12] of spine neck electrostatics, albeit with the assumption that only positive ions contribute to the overall current. This is an important effect which is in contrast to cable theory where the ion concentration is fixed to c_0_. Finally, the long and sustained synaptic input (Heaviside function) that leads to significant, long-term changes in spine head concentration and neck resistance is a condition that we used only to illustrate the slow-fast kinetics of the coarse-grained system of equations (11–12). Indeed, the kinetics of opening and closure of AMPA channels is rather of the order of few milliseconds [22]. To explore whether changes in concentration and neck resistance are already significant at this time scale during spontaneous synaptic activity, we used recent electrical recordings of voltage transients in spine heads and estimated the corresponding currents and changes in ion concentration for different putative spine geometries.

### Exploring the possible role of electrodiffusion with direct electrical recordings

Recently, we demonstrated the first direct measurements of spontaneous EPSP from spines using nanopipettes [8]. These recordings revealed voltage changes on a millisecond time scale with a fast rising phase (≈ 1 ms), followed by a slower decay phase (≈ 10 ms). A recent study measuring synaptic input currents revealed a similar time scale [15], which clearly indicates that the spine electrical transient is large and fast. These time-scales are much slower than the charging and discharging time constants of the spine head capacitor (tens of micro-seconds). Thus, it is most likely that the rising and decay phases correspond to the opening and closing kinetics of ion channels. We used the time course of this published data along with models of the rising and decay phase of ion channels [22] to explore the effect of electrodiffusion under different spine geometries. Although these recordings were made on spines with relatively long (>1 μm) necks, the large, fast, and spontaneous millisecond apart EPSPs could be used as a test input to explore the role of electrodiffusion on different spines geometries. We modeled the rising phase of synaptic conductance *g*^+^(*t*) with a sigmoidal function followed by a mono-exponential decay phase: 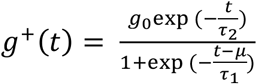 (Fig. 3a). To determine the conductance parameters for each EPSP, we used the following grid-search fitting procedure (Fig. 3a): Spine geometry is arbitrary fixed and for each synaptic event, we compute synaptic current (Eq. 10) and corresponding voltage time-course (Eq. 12) for a large range of conductance parameters [*g*_0_, *μ, τ*_1_, *τ*_2_]. The optimal set of parameters is then determined by minimizing the least-square distance between computed and measured voltage. We found a rapid opening kinetics (median *μ* = 0.52 ms and *τ*_1_ = 0.11 ms) followed by a slower decay (median *τ*_2_ = 3.95 ms) (Fig. 3a and Table 1). Given the estimated channel kinetics, we used the coarse grained electro-physiological model (equations (12–13)) to predict the kinetics of the ion concentration (Fig. 3b). We observed that the spontaneous variations of ion concentration were significant, particularly within spines with small head (reduced volume) and large necks (high current), leading to important reduction of neck resistance (Fig. 3c) and reversal potential (Fig. 3d). The kinetics of concentration variations was much slower than the EPSP time course, and for large EPSPs, we observed that the head concentration did not necessarily return to its resting state *c*_0_ before the arrival of a second EPSP. We thus predict that the concentration increase (i.e. resistance decrease) could result in a significant reduction in neck resistance during repeated synaptic stimulations (Fig. 4). This finding has one important implication in that, effective electrical resistance of the spine neck is fundamentally dynamics as it varies with synaptic activity and associated changes in concentration. Lacking other compensatory or homeostatic mechanisms, the increasing reduction in neck resistance during a train of EPSPs will effectively collapse the filtering effect of the neck resistance, leading to synaptic facilitation.

**Figure 3:**
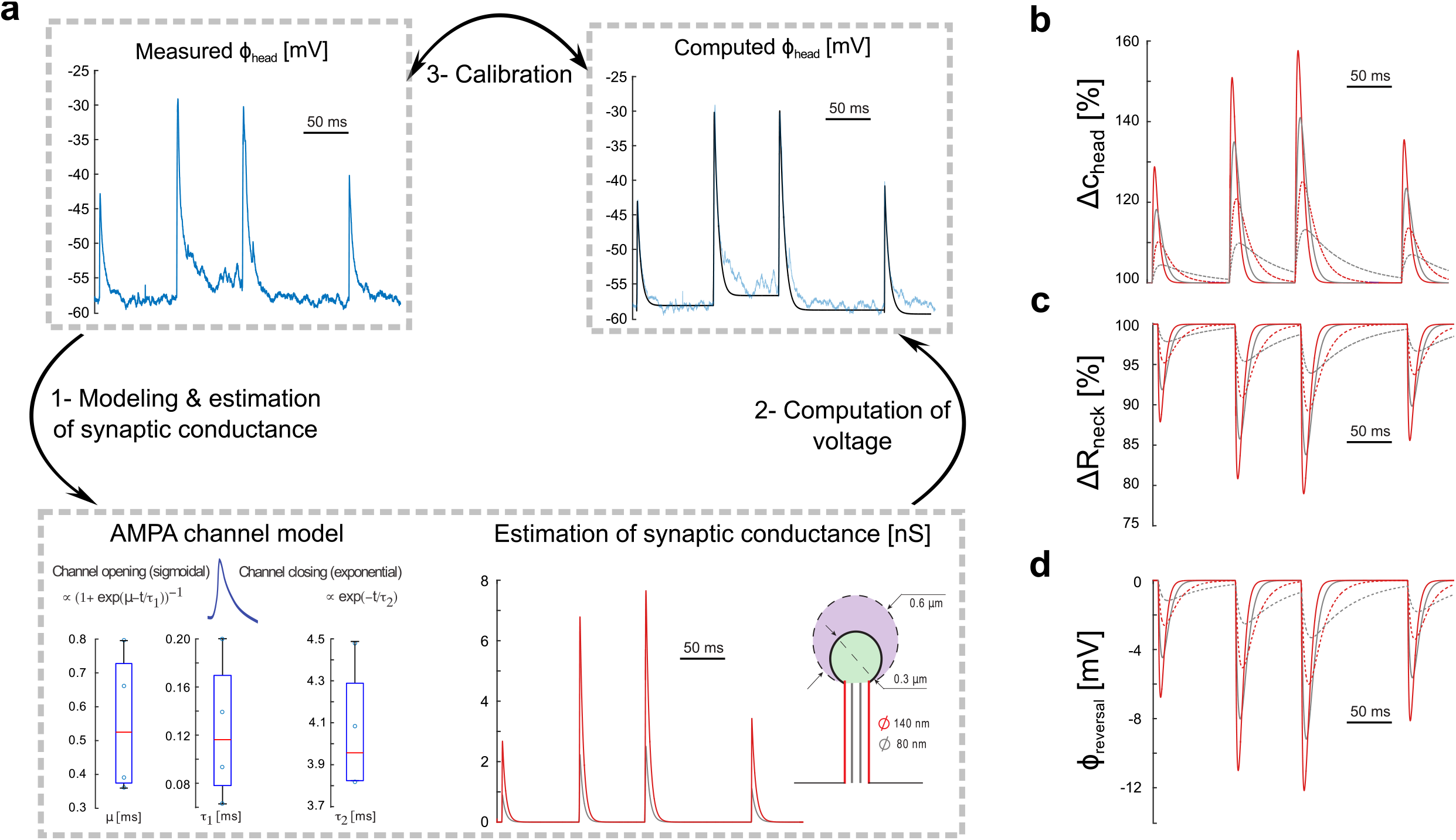
Experimental comparison. **a**) Estimating the synaptic conductance during spontaneous activity. 1- Voltage inside a single spine head during spontaneous activity, recently recorded using nano-pipettes (Jayant et al, Nat. Nano 2017). Conductance of AMPA receptors is modeled with sigmoidal opening (parameters μ and τ_1_) and single-exponential closure (parameter τ_2_). Boxplots show fitted parameters for the 4 EPSPs recorded experimentally, and optimal conductance is plotted for large (140 nm, red) and thin (80 nm, grey) spine necks. 2- To estimate optimal conductance parameters [g_0_, μ, τ_1_, τ_2_] for each EPSP, we performed a multi-dimensional grid-search algorithm where we computed the conductance g^+^(t), synaptic current (Eq. 10) and voltage (Eq. 12) for a large range of parameters. 3- Calibration of best conductance parameters was performed by minimizing the distance between the computed and measured EPSPs. **b**) Dynamics of concentration, **c**) Neck resistance and d) Reversal potential as reflected by coarse-grained electro-diffusion model during spontaneous spine activity in different spine geometries (large (R = 600 nm, dashed line) and small (R= 300 nm, solid lines) head; large (140 nm, red) and thin (80 nm, grey) necks). Notice that spines with small head and large necks will see the maximal effect of concentration change on neck resistance.

**Figure 4:**
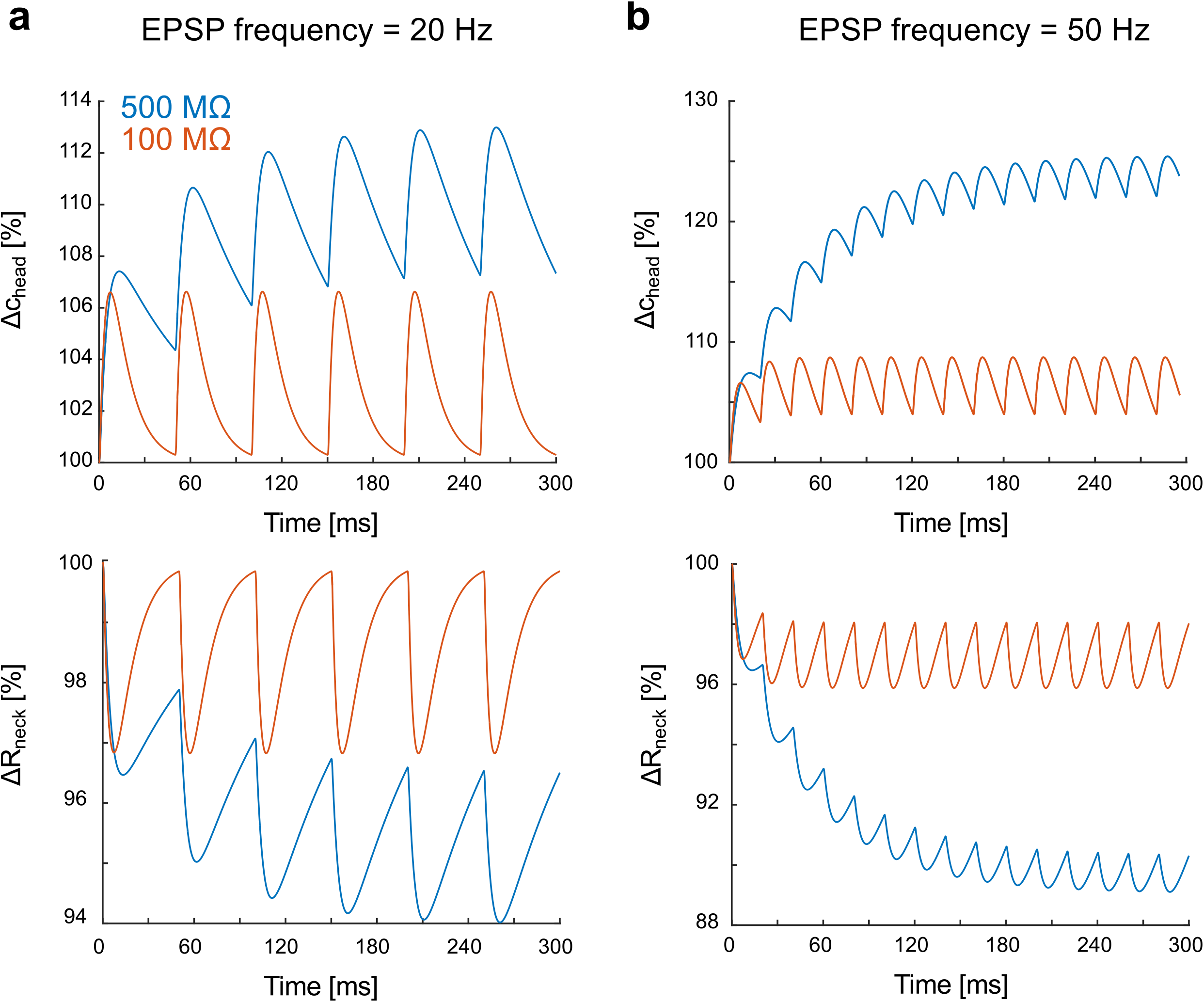
Incremental increase of ion concentration and associated neck resistance decrease during high-frequency synaptic stimulation. **a**) Relative variations of ion concentration in the spine head (R= 300 nm) and associated neck resistance for a 20 Hz synaptic stimulation (geometrical neck resistances (i.e. at concentration c_0_): 500 MΩ (blue) and 100 MΩ (red)). The kinetics of the synaptic conductance is 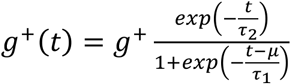 with g^+^ = 2 nS, μ = 0.55 ms, τ_1_ = 0.12 ms and τ_2_ = 4 ms. **b**) Relative variations for 50 Hz synaptic stimulation. We do point out that in contrast to Fig.3, we did not impose a voltage amplitude or profile in the spine head in the above analysis, but rather subjected the spine to trains of given synaptic inputs at different frequencies. The difference is that the trend in dynamic modulation is reversed: low resistance neck resistance spines show less modulation while high neck resistance spines show a more dynamic change.

## DISCUSSION

The spine neck is an important barrier to ion diffusion during synaptic transmission, but its role in electrically shaping EPSPs has remained controversial due to the lack and difficulty in performing precise experimental measurements from dendritic spines. This has led to an incomplete and often contradictory understanding of spine electrical properties [7, 8, 10, 11, 14, 17, 41, 42]. One way to overcome this handicap is with accurate biophysical models. This is normally achieved with cable theory modeling, which is widely used in simulations of neuronal physiology. However, cable equations neglect the role of electrodiffusion (i.e. electric field effect on ion gradients), which can become appreciable in small neuronal compartments such as spines [34]. Here, we use instead an electrodiffusion framework and model the electrostatics inside the spine during synaptic stimulation. Following previous efforts using non electroneutral conditions [34] or numerical simulations [12], here, using singular perturbation theory of the PNP equations to model kinetics, we derive a coarse-grained model that fully captures the coupled dynamics of ion concentration and potential inside the spine head. Specifically, we find that (i) diffusion and electrodiffusion jointly govern the kinetics of spine excitatory post-synaptic potentials (EPSPs); (ii) the spine geometry plays a key role in shaping the EPSP time course; and, (iii) that the current-voltage relationship across the spine-neck is non-linear, which results in the neck resistance varying as a function of ion concentration. We briefly discuss the functional implications of the above findings.

### Effect of electrodiffusion in dendritic spines

Using a coupled slow-fast dynamical system analysis with a Heaviside step input waveform, we find that the voltage transient can be divided into an *electrostatic phase* - lasting a few hundred microseconds; and a *diffusion* phase - lasting several milliseconds. This result shows that the electrical voltage rises very fast due to fast charging of the spine head (due to a low membrane capacitance), with a steady-state value determined by *I_syn_* × *R_neck_*, while the diffusion of ions begins to occur only a few milliseconds later. But, if the spine EPSP is sufficiently high, this adds a significant electromotive force on the ions driving them out of the neck, contributing to a fast and large synaptic current. After few microseconds (Fig. 2g), head capacitor is charged and neck current equilibrates with synaptic current. Amplitude of synaptic current is proportional to receptors’conductance but also to the difference between head potential and reversal potential, a difference which decreases as ion concentration builds-up in the spine head (Eq. 10). Because of this, synaptic current also depends on spine geometry and neck resistance. Finally, we find that the effect of spine geometry on spine and neck resistances and on the voltage gradients across the spine neck can be further exaggerated by local membrane curvature and constriction of the neck, and also by the reduction of the apparent cross-section due to crowding with organelles such as spine apparatus [43]. These effects might significantly increase neck resistance as it is inversely proportional to neck (apparent) cross-section (Eq. 54 in SI).

### Regulation of spine neck resistance by ionic concentration

As expected, we find that the rapid voltage increase during the opening of AMPA receptors drives electrical currents through the spine neck. Neck current here corresponds to an exchange between positive ions flowing from the spine head to the dendrite with negative ions flowing from the dendrite to the spine head. After few microseconds, when the head capacitor is charged, synaptic and neck currents equalize. But, as synaptic current only involves positive ions whereas neck current results from an exchange between positive and negative ions, a fraction of entering sodium ions remains inside the spine, and their relatively slow diffusion through the neck enables a gradual accumulation inside the head nanodomain during receptors opening. At the same time, an inward neck current of negative ions maintains the overall electro-neutrality.

Our model predicts that the currents associated with recorded EPSPs change ion concentration inside the spine by up to 50%. This agrees with recent experiments using fluorescent sodium indicators that reported up to 10 mM concentration increase following the AMPA receptors opening during EPSPs [25] and large synaptic conductance’s ranging between 2-8nS measured from spines [15]. It is important to point out that we made two major assumptions when modeling ion dynamics: First, we neglected the effect of potassium ions and calcium entering the spine neck, which could further change the electrostatic landscape. For example, potassium ions could enter the spine neck from the dendrite in the event of SK channel dependent shunting [44]. This current could counter the decrease in spine neck resistance due to the increased sodium concentration and thus tune the net synaptic current. The second important assumption of our model is that negative charges were only accounted for by diffusive chloride ions, whereas it is known that negative charges are also partly accounted for by less mobile proteins [45]. The slower effective diffusion of negative charges would increase the neck resistance (Eq. 8 and 9), and reduce the inward negative current. This could down-regulate changes in concentrations and electro-diffusion effects on reversal potential and neck resistance. But, regardless of the effects of other ions and motility, the sodium concentration build-up should then lead to a reduced electrical resistance through the neck. Moreover, as diffusive extrusion of accumulated ions from the spine head is relatively slow (ten’s to hundreds of milliseconds), we predict that high-frequency synaptic input will lead to a pronounced decrease in neck resistance, as the rate of ion concentration buildup in the spine nanodomain will exceed the rate of diffusion through the neck. Our analysis thus reveals a simple, yet new, physiological mechanism of post-synaptic facilitation at millisecond time scale.

To finish, we speculate that part of the reason that spine neck resistance measurements have been at odds with each other could be due to the enhanced ion concentration in the spine which dynamically changes the resistance. Therefore, in addition to the voltage dynamics in the spine head, the net current through the neck would be influenced by the net ion concentration inside the spine nanodomain, and care must be taken to measure those in order to properly interpret the electrical function of spines. This is something which is not of purely academic interest, as the extend to which spines implement electrical compartments and shape the kinetics and amplitudes of EPSPs is of fundamental importance to neuroscience, as they serve to mediate most excitatory transmission in the vertebrate central nervous system.

## Supplementary information (SI) accompanies this manuscript

SI includes detailed mathematical derivations of the equations presented here in this manuscript

## Acknowledgements

This work was supported by the NIMH (R01MH101218, R01MH100561) and the NINDS (R01NS110422). This material is also based upon work supported by, or in part by, the U. S. Army Research Laboratory and the U. S. Army Research Office under contract number W911NF-12-1-0594 (MURI). T.L. was partly supported by the Fondation pour la Recherche Médicale and the Philippe foundation. K.J was partly supported by the Kavli Institute of Brain Science at Columbia.

## Competing Financial Interest

Authors declare no competing financial interests pertaining to this study.

## Author Contributions

T.L. and R.Y. conceived of the project. T.L performed the modeling and analysis. K.J assisted with model development and analysis. T.L and K.J wrote the manuscript. R.Y assembled and directed the team, provided guidance, funding, and edited the manuscript.

